# Activation of pancreatic β-cell genes by multiplex epigenetic CRISPR-editing

**DOI:** 10.1101/2020.07.24.214544

**Authors:** Gimenez Carla Alejandra, Curti Lucia, Hyon Sung Ho, Grosembacher Luis, Ross Pablo Juan, Pereyra-Bonnet Federico

## Abstract

CRISPR-based systems for epigenetic editing are promising molecular tools that could be harnessed for directed differentiation of pluripotent stem cells. We used the CRISPR/dCas9-VP160, CRISPR/dCas9-TET1 and CRISPR/dCas9-P300 systems for multiplex epigenetic editing and activation of human beta pancreatic genes (*PDX1, NEUROG3, PAX4* and *INS*). The CRISPR/dCas9-P300 system was the most effective at activating genes with reduced number of sgRNA. Using small number of sgRNA per gene was important to induce multiplex gene activation. Combined activation of transcription factors (TFs) involved in beta cell development resulted in *INS* gene expression; in which sequential TFs activation was more effective than simultaneous activation. Full CRISPR RNA-based delivery system was able to activate all targeted genes. Overall, this study shows the utility of CRISPR tools for epigenetic editing and directed cellular differentiation.

## INTRODUCTION

Type 1 diabetes (T1D) is characterized by the loss of glucose homeostasis due to an autoimmune destruction of the insulin-producing cells in the pancreas. The most commonly used treatment for T1D is the exogenous administration of insulin; however, treatments that could provide superior control of blood glucose are needed.^1,2^ Cellular replacement therapies based on in vitro differentiated induced pluripotent stem cells (iPSCs) may offer a novel approach by enabling fine regulation of glucose homeostasis, which in turn will reduce the need for constant monitoring and also prevent long term micro and macrovascular complications.^3–7^

In vitro differentiation methods mimic the processes triggered during embryonic development towards a certain cell type, tissue or organ exposing cells to different signaling molecules or by exogenous gene overexpression.^8,9^ However, differentiation and activation of the critical transcriptor factor (TF) responsible for development and achieving a matured phenotype is often incomplete.^10,11^ Therefore, tools that allow on-demand direct endogenous gene activation could make reprogramming methods towards specific destinations more effective and efficient.

Recently, new CRISPR-Cas9-based approaches to regulate gene expression have been developed. These approaches known as CRISPR activator systems (CRISPRa) use a mutated “dead” Cas9 (dCas9) protein without endonuclease activity.^12–14^ The dCas9 is fused with an epigenetic effector domain to allow for targeted epigenetic editing. The high level of accuracy of the CRISPR system with the addition of effector domains allows introducing epigenetic modifications at specific location within the genome, a process known as epigenetic editing.^15,16^ One of the most commonly used effector domains is the transcriptional activation domain of the VP16 human papillomavirus oligomers with different copy numbers (e.g. VP64, VP160)^12,17,18^, similarly “epigenetic erasers” such as TET1^19^, and “epigenetic writers” such as the core of the p300 histone acetylation protein have been used in this system.^20^

Our group and others found that using the CRISPR/dCas9 platform with VP160 effector and several sgRNA at the same time is associated with a synergistic stimulus for gene activation^17,18,21^. On the contrary, the use of CRISPR/dCas9 associated with p300 can be used efficiently with a single sgRNA.^20^ While CRISPR gene editing was extensively used as a multiplex platform for modifying several genes at the same time,^22–32^ only few reports showed the multiplexing capacity of CRISPRa platform,^17,18,33^ and none of them in a significant number of genes corresponding to the same cellular lineage.

In this study, the goal was to demonstrate the feasibility of using the CRISPR/dCas9 platform with different effectors to modify epigenetic states and activate multiple genes related to pancreatic beta-cell development (*PDX1, NEUROG3, PAX4* and *INS*). We compared and optimized multiple sgRNAs, and their combinations, for achieving highest effects, as well as tested different strategies for multiplexed epigenetic editing and directed cell differentiation.

## METHODS AND MATERIALS

### Ethics Statement

The protocols were approved by the Institutional Ethics Committee of the Hospital Italiano, Argentina (Res 1672 and 2251).

### sgRNA design

The *NEUROG3* and *PAX4* sgRNAs were designed using the CRISPR Design Tool (Feng Zhang Lab, MIT). The algorithm used by this program is based on a previously described specificity analysis^34^. The *INS* sgRNAs and the *PDX1* sgRNAs were previously reported^21,33^

### Plasmids and sgRNAs preparation

All plasmid vectors used in this study were obtained from Addgene (plasmids #47108, #48226, #61357, #83889 and #82559 respectively).^18–20,35^ The sgRNAs oligonucleotides containing the target sequences were cloned as described by Ran and colleagues.^23^ Chemically modified synthetic sgRNA (2’-O-methyl analogs and 3’ phosphorothioate internucleotide linkages) were purchased from Synthego (CA, USA). Sequences of oligonucleotide used for sgRNA cloning and synthetic sgRNAs are provided in Supplementary Tables S1 and S2.

### Cell culture and transfection

HEK293T transfections were performed using Lipofectamine 2000 (Invitrogen; Carlsbad, California) using a 1:1 DNA/reagent ratio. For individual gene activation experiments, the dCas9 plasmids were transfected at a mass ratio of 1:1 to either the individual sgRNA expression plasmids or the identical amount of sgRNA expression plasmid consisting of a mixture of equal amounts of each sgRNAs. For multiplex gene activation experiments see Supplementary Table S3. Control group cells were transfected with the dCas9 plasmids and empty sgRNA expression plasmids at a mass ratio of 1:1. hIPS cells were cultured in Essential 8™ Medium with E8 Supplement (Gibco, USA) and transfected using Lipofectamine™ MessengerMAX™ Transfection Reagent (Invitrogen, USA) according to the manufacturer’s instructions.

For testing different culture medium conditions, cells were cultured with DMEM medium (90%DMEM, 10%FBS and 1% antibiotics) for 4 days and then continued on DMEM medium or changed to TM medium [40 ng/mL bFGF, 20% xeno-free serum replacement (Invitrogen), 2 mM glutamine, 0.1 mM nonessential amino acids, 0.1 mM β-mercaptoethanol, and 2% antibiotics in DMEM/F12 Knockout (Invitrogen)] with either 50ng/ml exendine-4 (Ex-4, Baxter, Indiana, USA) or 25 ng/ml Trichostatin A (TSA-) and 1 µM 5-azacytidine. Immunocytochemistry analysis was performed on day 10 of culture. Cell transfection scheme is detailed on Figure 5A.

### HEK293-LV-dCas9-P300 and hiPS dCas9-P300 cell line production

For production of dCas9-P300 lentiviral particles, envelope and capsid plasmids obtained from Addgene (#83889, #12260 and #12259) were used. HEK293T cells were transfected at 4:3:1 dCas9-P300 lentiviral plasmid: capsid: envelope ratio. Target cells were infected by spinfection protocol as described by Shalem and colleagues.^26^ Next, fresh DMEM medium was placed onto plates and incubated at 37 °C overnight. For clone selection puromycin selection was performed for one week at 0.5-1 µg/ml concentration. Clones were selected, expanded and tested for dCas9-P300 expression by qPCR.

### Gene and protein expression analysis

Total RNA was isolated with RNeasy Mini Kit (Qiagen). For RT-PCR and qPCR, the RNA was reverse-transcribed using ImProm-II™ Reverse Transcriptase (Promega, Massachusetts, USA). Specific intron-spanning primers were used in the PCR and qPCR (Supplementary Table S4). The qPCR was performed using KAPA SYBR® FAST qPCR Kit Master Mix (2X) Universal (Kapa Biosystems; Massachusetts, USA). Protein expression at day 10 post transfection was analyzed by immunocytochemistry using the following antibodies and dilutions: PDX1 1:100 (134150, Abcam), NEUROG3 1:200 (HPA039785), PAX4 1:100 (AV32064, Abcam) and INS 1:100 (181547, Abcam).

### ChIP-qPCR analysis

HEK293T cells were collected 4 days after co-transfection of dCas9/P300 and the most efficient sgRNAs per gene (5 sgRNAs in total). ChIP was performed as described by Lin and colleagues.^36^ Primers used for the qPCR are listed in Supplementary Table S4. The percent input method was used to analyze the data.^37^

### DNA methylation analysis

Genomic DNA from CRISPR-on HEK293T cells and control cells were purified using a AllPrep DNA/RNA/Protein Mini Kit (Qiagen). Then, the DNA was treated with KruO4 for 5hmC recognition according to a previous report^38^ and treated using a EpiTect Bisulfite Kit (Qiagen) as described.^21^ The post-bisulfite promoter region of the human *INS* gene (NCBI ID 3630) was amplified by PCR using specific primers (Supplementary Table S4).

### *In vitro* transcription (IVT)

DNA template was amplified from dCas9-P300 expressing plasmid (Addgene #48226) using specific primers for IVT reaction (Supplementary Table S4). The conversion of DNA to RNA was carried out using the MegaScript® SP6 kit (AM1330, Ambion, Invitrogen), according to the manufacturer’s instructions with minor modifications. Briefly, the protocol was modified by adding a cap analog (3’-O-Me-m7G) (5 ‘) ppp (5’) G RNA Cap Structure Analog, (AM8045, Thermo Fisher), in a 4: 1 ratio with the GTP nucleotide. Next, a treatment with Antarctic Phosphatase (M0289S, New England Biolabs) was performed for 30 minutes.^39^ The resulting RNA was purified by the MegaClear® kit (AM1908, Ambion, Invitrogen), according to the manufacturer’s instructions. For visualization, synthetic RNAs were run on a gel under denaturing conditions.

### Statistical analysis

All of the data are presented as mean ± SEM and represent 2-4 biological replicates with each “n” value and test choice reported in the figure legends. Data from fold change gene expression experiments were not normally distributed and were log transformed for the statistical analysis. Statistical analyses were performed with GraphPad Prism 5.0 software. A value of p<0.05 was considered significant. Results are represented as *p<0.05.

Supplementary information is available at The CRISPR Journal website

## RESULTS

### Capacity of different CRISPR/dCas9 systems to activate pancreatic genes

To induce transcriptional activation of *PDX1, NEUROG3, PAX4* and *INS* genes, we tested different CRISPR-dCas9 activation systems, including the dCas9-VP160, dCas9-TET1, and dCas9-P300 (Fig. 1A). Initially, we used a combination of 4-5 sgRNAs targeting the promoter region of each gene (Fig. 1B, Supplementary Table S1). RT-qPCR was performed 4 days after transfection to assess activation of the targeted genes (Fig. 2A). dCas9-P300 was effective at activating transcription of *all tested genes*. The specific activity of dCas9-P300 was corroborated by increased levels of histone acetylation at targeted regions of *NEUROG3* and *PAX4* (Fig. 2B). dCas9-VP160 was effective at activating *NEUROG3, PAX4* and *INS*., but did not efficiently induce *PDX1*. Analysis of DNA methylation levels at the *INS* promoter confirmed demethylation induced by dCas9-TET1 system (Fig. 2C); however, we did not observe transcriptional activation for any of the targeted genes except for *PDX1* (Fig. 1A). Transfection of all systems simultaneously, did not improve transcriptional activation, indicating a lack of synergistic effects between the three tested systems (Fig. 2A). Overall, dCas9-P300 and dCas9-VP160 were able to activate gene expression of pancreatic genes in HEK293 cells.

**Figure 1.**
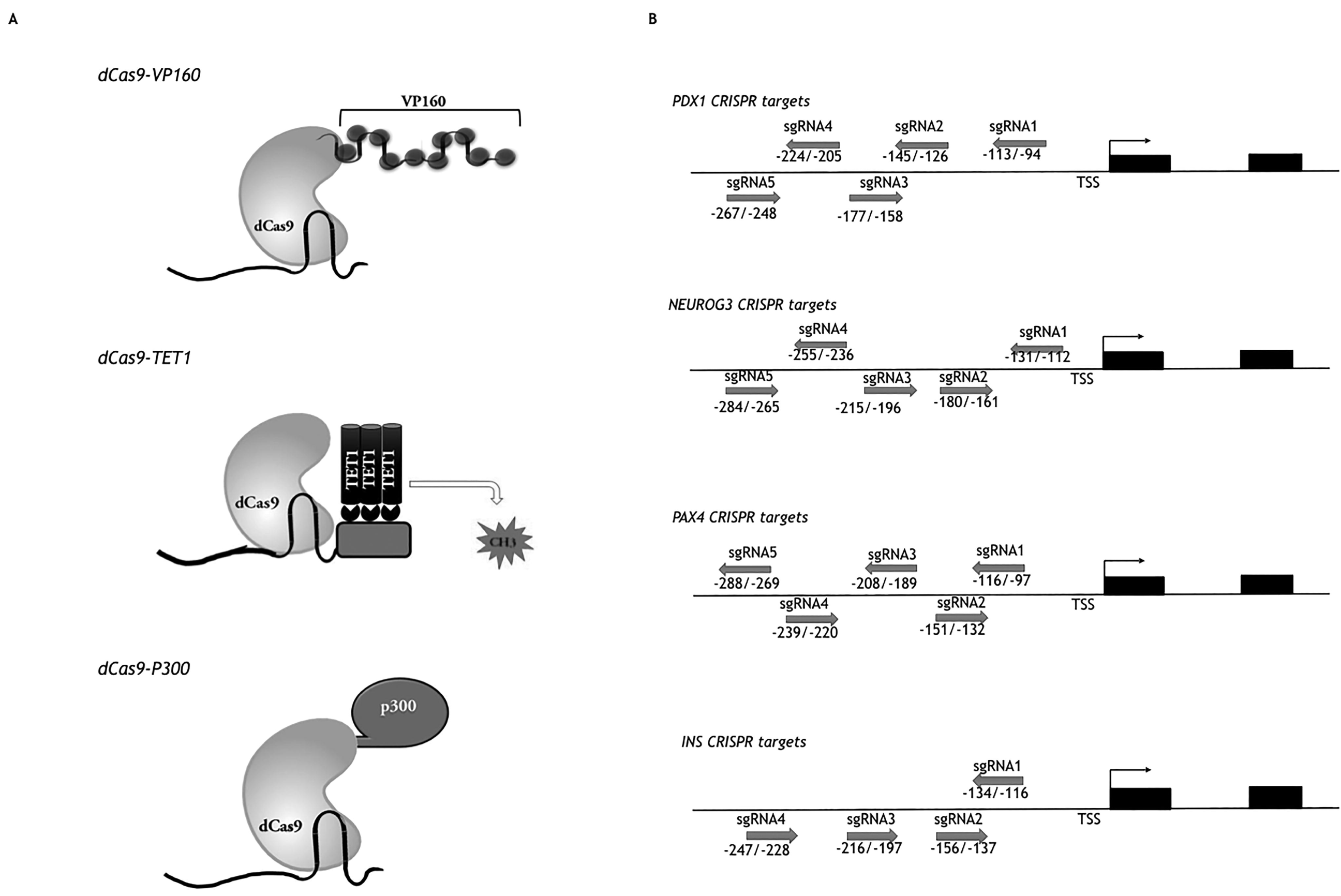
CRISPR activation systems and sgRNAs targets tested. (A) CRISPR activation systems schemes: dCas9-VP160 for transcription machinery attraction, dCas9-TET1 for targeted DNA demethylation, and dCas9-P300 for histone acetylation in H3K27. (B) sgRNAs target positions on *PDX1, NEUROG3, PAX4* and *INS* proximal promoter. Numbers refers to sgRNA nucleotide position respect to transcription start site (TSS) according to NCBI database (Gene ID for *PDX1:*3651, *NEUROG3:*50674, *PAX4:*5078 and *INS*:3630).

**Figure 2.**
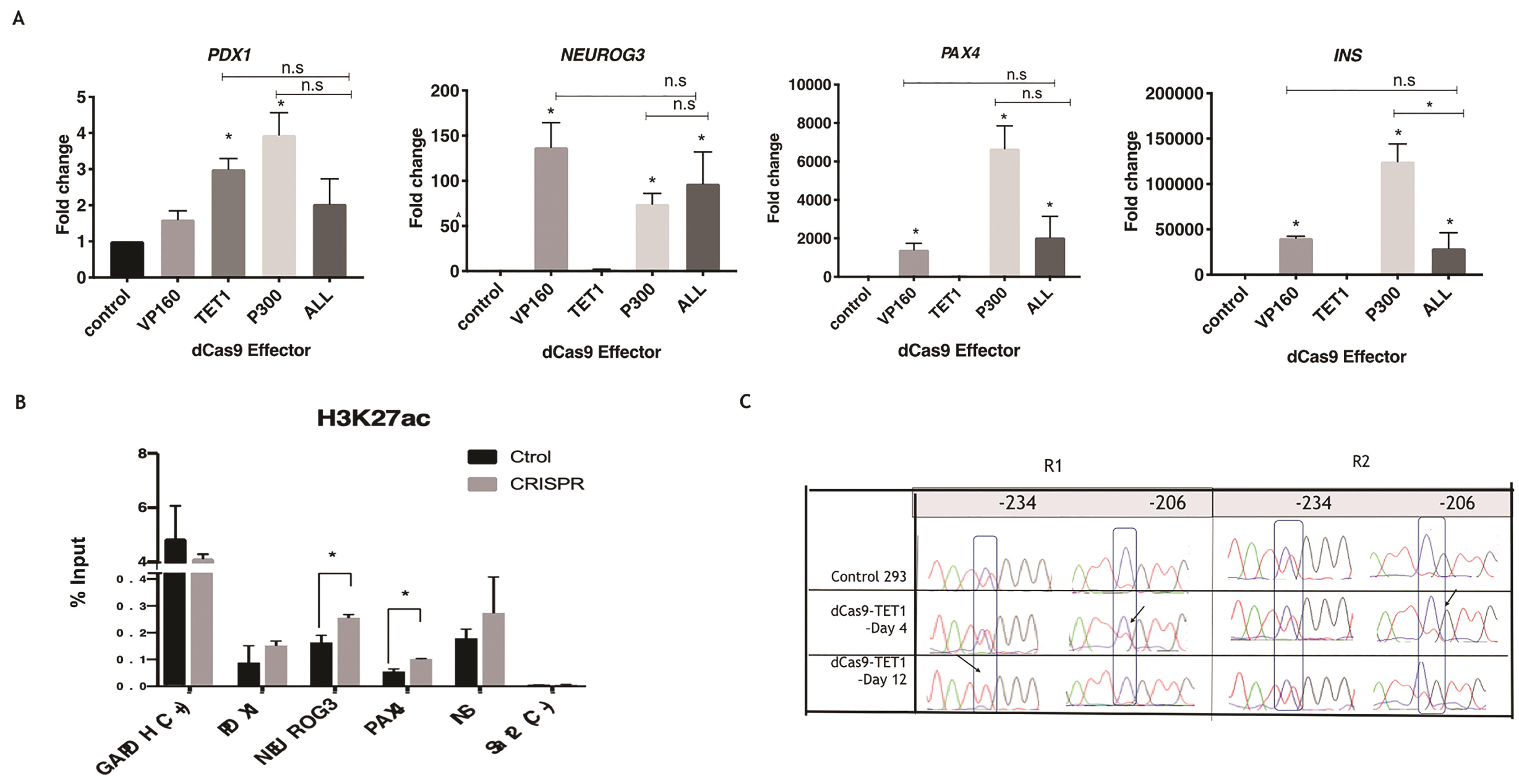
Multiplex activation of endogenous lineage pancreatic genes with different CRISPR dCas9 activation systems. (A) RT-qPCR gene expression analysis on multiplex gene activation for each CRISPR system. Gene expression levels of *PDX1, NEUROG3, PAX4* and *INS* in the human HEK293 line when different dCas9 Effector systems are transfected for activation (VP160, TET1 or P300) separately and together. For all cases, the tests were performed using all the sgRNAs for all 4 genes tested at the same time. Bars show fold change respect control group + SEM and One way ANOVA-Turkey test was used. N = 3 biological replicates for all groups with the exception of TET1, which has 2 biological replicates. Unless otherwise marked, “*” shows significance respect control group. (B) H3K27ac levels for dCas9-P300 multiplex gene activation, t-student test was used (n= 2 biological replicates). (C) *INS* promoter methylation level in HEK293 transfected with dCas9-TET1 system at day 4 and 12. CpG -234 y -206 respect to TSS are shown. The table shows the methylated cytosine studied in the control (blue) and how it is demethylated and converted to thymine post-PCR bisulfite (red). The arrows indicate the increase in thymine level and the decrease in cytosine level.

### Single and combinatorial use of gRNA for CRISPR-based transcriptional activation of pancreatic genes

To optimize transcriptional activation, we determined the effect of each individual sgRNAs on driving dCas9-VP160 and dCas9-P300 gene activation. The dCas9-VP160 system showed minimal transcriptional activation when single sgRNA were used, while significant activation levels were observed when all sgRNAs were combined (Fig 3A). On the other hand, dCas9-P300 achieved maximal levels of P*DX1, NEUROG3* and *PAX4* transcriptional activation with unique individual sgRNAs (Fig 3B). Combining 2-3 of the most efficient sgRNA identified for dCas9-P300 did not achieve improved gene activation; except for *INS*, which required a combination of the two most efficient sgRNAs to obtain maximal transcriptional activation (Fig. 3C). Furthermore, using immunostaining we determined that transcriptional activation corresponded with increased protein expression in HEK293 cells (Fig 3C).

**Figure 3.**
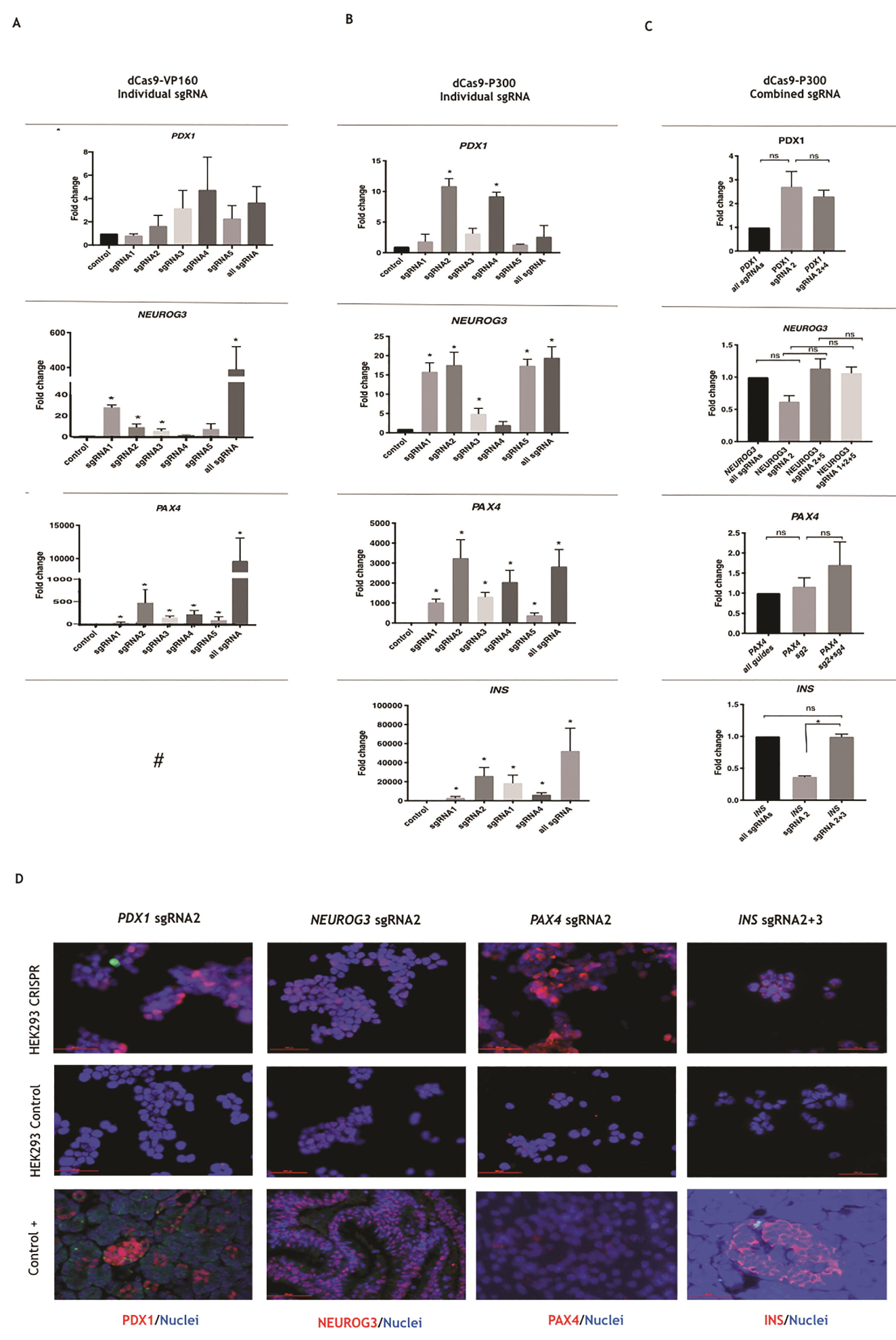
Single and combined sgRNAs performance on the CRISPR dCas9 activation systems. (A, B) RT-qPCR gene expression analysis of *PDX1, NEUROG3, PAX4* and *INS* in the human HEK293 line when individual and combined sgRNA are transfected with the dCas9-VP160 and dCas9-P300 systems (n=3-4). (C) Combination of the most efficient sgRNAs vs *All sgRNA* group for dCas9-P300 (n=2). For all cases, the tests were performed for each gene individually. For the “all sgRNAs” group (all sgRNAs per gene) the total mass was divided by the number of different sgRNAs to be used. Bars show fold change respect control group + SEM and One way ANOVA-Turkey test was used. Unless otherwise marked, “*” shows significance respect control group. “n.s”: not significant. #: tested before by our group.^8^ (D) Inmunocytochemistry for individual gene activation experiments in dCas9-P300 expressing cell line using the most efficient guides at day 10 post transfection.

### Multiplex pancreatic gene activation in HEK293 cells

Given that dCas9-P300 system was efficient at activating gene expression with only 1 or 2 sgRNAs, we tested the possibility of activating multiple genes simultaneously. For this experiment, we used a cell line that constitutively expresses dCas9-P300 (HEK293-LV-dCas9-P300), and therefore only sgRNAs needed to be transfected (Supplementary Fig. S1). RT-qPCR at day 4 post-transfection revealed increased gene expression for all targeted genes, in contrast to the same experiment using all sgRNAs, in which only 3 out of 4 genes were activated (Fig. 4A). These results show that that multiplex gene activation can be achieved using a minimal number of optimized sgRNA combined with dCas9-P300.

**Figure 4.**
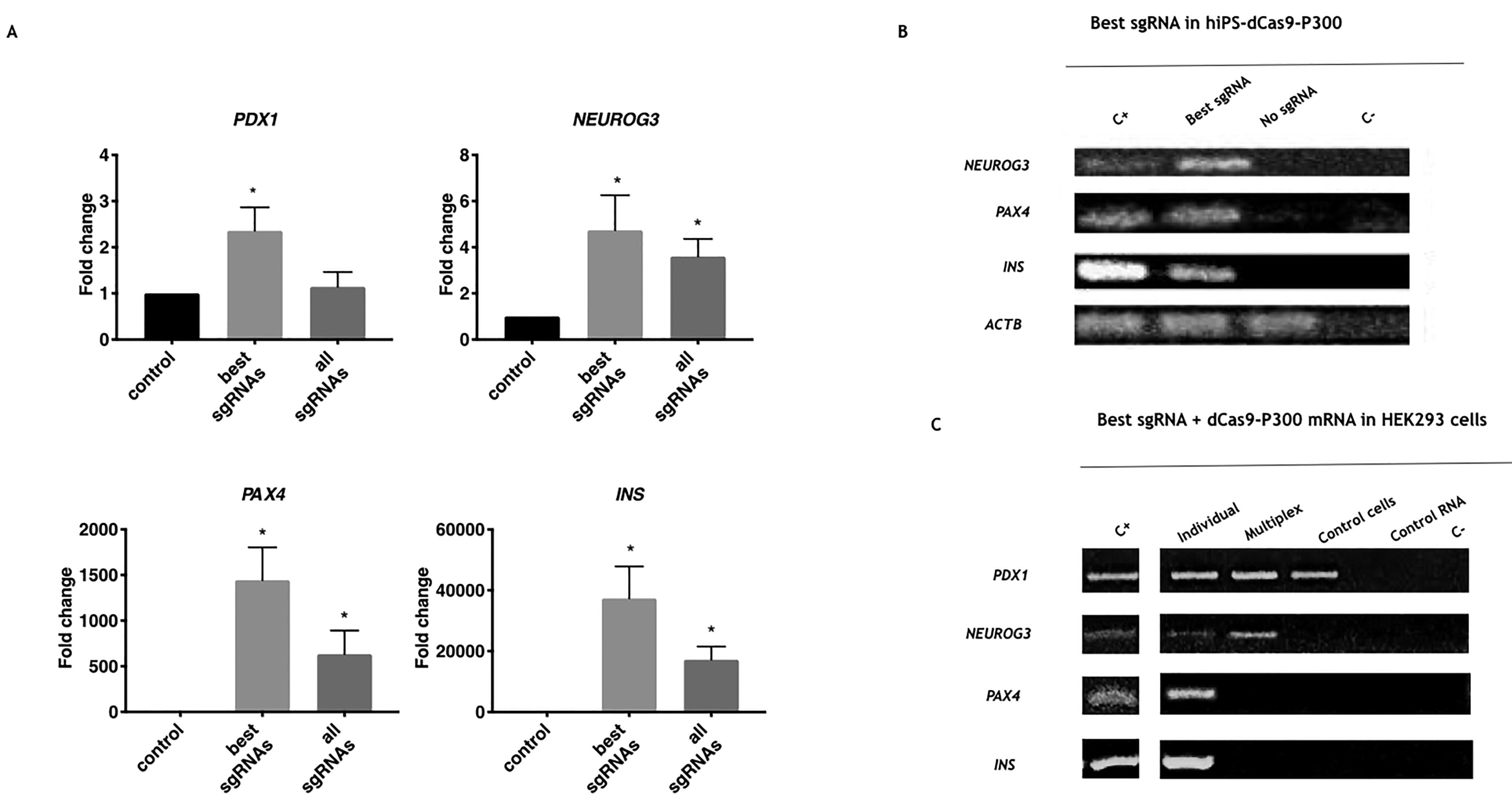
Multiplex activation of endogenous lineage pancreatic genes. (A) RT-qPCR gene expression comparasion analysis of the most efficient sgRNAs vs all sgRNAs in mutiplex activation gene experiments (n=4). For all analysis, bars show fold change respect control group + SEM and One way ANOVA-Turkey test was used. “*” shows significance respect control group. (B) RT-PCR in hIPS dCas9-P300 cell line transfected with the most efficient synthetic sgRNAs for *INS, NEUROG3, PAX4*. (C) RT-PCR beta pancreatic gene expression analysis in ordinary HEK293 line transfected with dCas9-P300 mature RNA and the most efficient synthetic sgRNAs. “C+” is human pancreas cDNA; “Individual genes *INS, PAX4, NEUROG3, PDX1*” are cell groups transfected with the corresponding most efficient sgRNA of each gene; “Multiplex” is the cell group transfected with the most efficient sgRNAs from all genes; “Ctrol cells” is the group without any sgRNA transfected; “Ctrol RNA” is RNA treated by DNAse before cDNA convertion and “C-” is PCR mix control.

In order to develop a clinically friendly system for gene activation, we tested the possibility of replacing sgRNA plasmid/lentiviral delivery by synthetic RNA molecules, in a relevant cell type (iPSCs). Transfection of synthetic RNA guides into hiPSC overexpressing dCas9-P300 resulted in activation of all target genes, demonstrating the effectiveness and robustness of the strategy in multiple cell types (Fig.4B and Supplementary Fig. S2). Finally, we combined synthetic sgRNAs with dCas9-P300 mRNA to generate an entirely RNA-based method for individual and multiplex gene activation. Delivery of the RNA-based CRISPRa system in HEK293 cells resulted in transcriptional activation of all targeted genes when individually targeted, but not in multiplex activation attempts (Fig. 4C and Supplementary Fig. 3).

Given that PDX1, NEUROG3, and PAX4 are transcription factors (TFs) involved in pancreatic beta cell differentiation, we tested several activation schemes, including single TFs, simultaneous activation of all three TFs, and sequential activation of the three TFs, for the ability of inducing *INS* expression. We also tested different culture medium conditions, including basal culture medium, TM culture medium plus exenatide (a common chemical used in beta pancreatic differentiation) and TM culture medium supplemented with 5 aza-cytidine and TSA (as epigenetic barriers mediated by chromatin modifiers can also hinder lineage reprogramming)^40^ (Fig. 5A). Using immunostaining 10d-post transfection, we observed cells with INS+ signal in their cytoplasm. When cells were cultured in basal DMEM medium, we observed INS signal in NEUROG3 sgRNA transfected cells and in the sequential activation group. For TM culture medium plus exenatide, signal was observed in every group, although at different intensities. For TM culture medium plus 5 aza-cytidine and TSA, signal was seen in *PDX1* sgRNA, *NEUROG3* sgRNA and the sequential group. No INS^+^ cells were observed in control groups (Fig. 5B). As a result, the best individual sgRNA activation strategy was achieved when we used NEUROG3 independent of the media and simultaneous or sequential activation. The use of exenatide showed INS expression in all groups which indicates that exenatide supports the best conditions for differentiation; and sequential use of the CRISPRa stimulation resulted in INS expression in all tested culture conditions.

**Figure 5.**
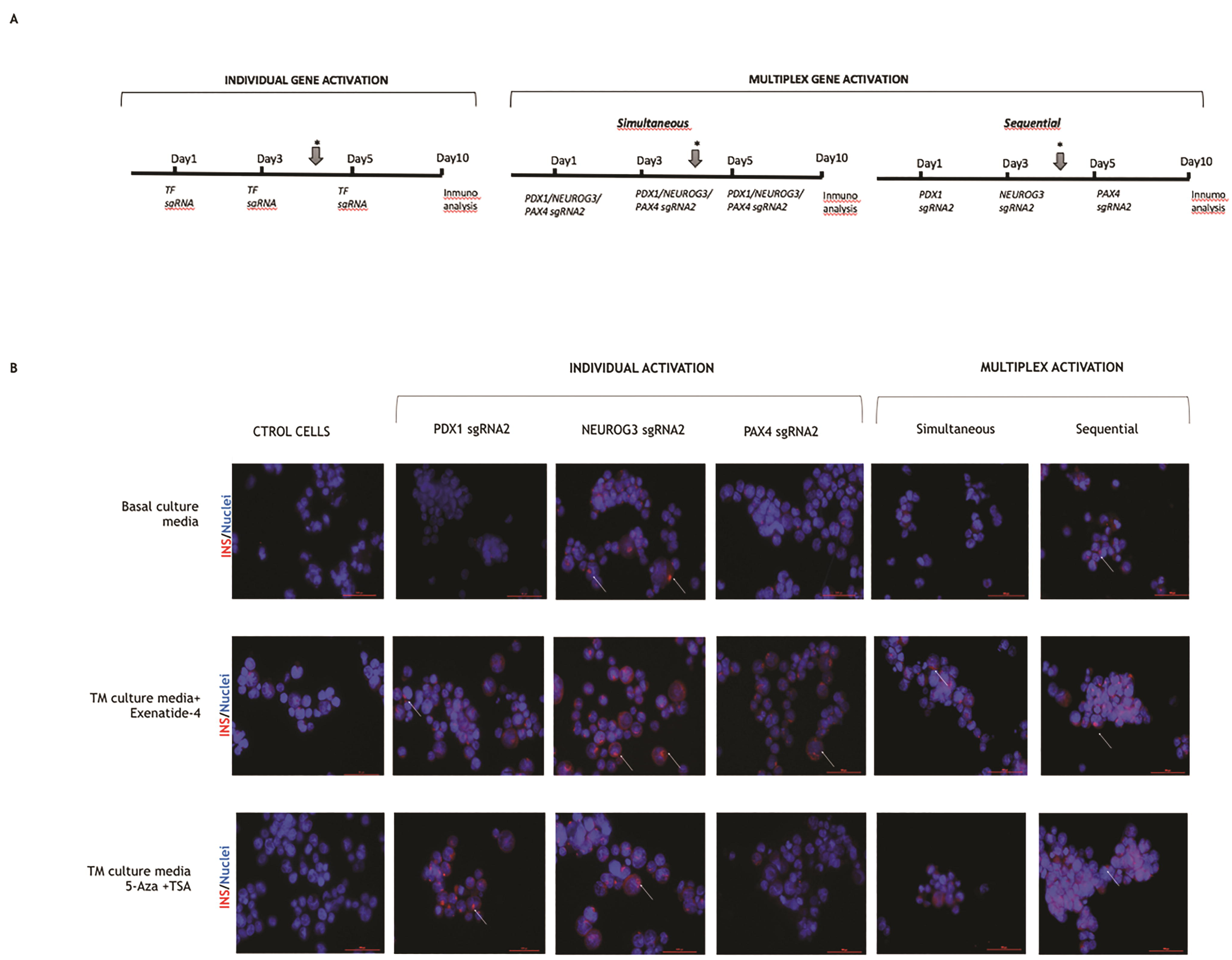
Protein expression for individual, simultaneous and sequential transcription factor CRISPR activation in different media. (A) Transfections scheme for individual, simultaneous or sequential gene activations in different culture media. Arrow represent medium change day. (B) Immunocytochemistry at day 10 post transfection for INS in different TF CRISPR activation groups and cell culture medium (40X). Arrows show INS signal cumulus.

## DISCUSSION

In this study, we tested several parameters to determinate which are the best CRISPRa tools and conditions to use in cellular protocols aimed at inducing β-cell differentiation *in vitro* minimizing cost and time.

The dCas9-P300 and dCas9-VP160 systems were the most effective at activating β-cell genes *in vitro*. The dCas9-TET1 system did not significantly activate most of our target genes, even though partial DNA demethylation was corroborated at the *INS* gene. This result was not surprising considering that demethylation the promoter regions does not imply gene activation *per se*.^41^ Contrary to previous reports on CRISPR systems for gene silencing,^42^ the simultaneous use of different type of CRISPRa systems did not have a synergistic effect.

From all CRISPRa systems, dCas9-P300 was most effective at activating β-cell genes with reduced number of gRNAs. In concordance with previous work, we observed that dCas9-VP160 system requires several sgRNAs per gene to achieve a significant activation effect,^17,18,21^ while dCas9-P300 reaches similar activation levels using only one sgRNA.^20^ The different efficiencies could be attributed to the capacity of dCas9-P300 to induce a direct epigenetic modification, by acetylation of H3K27, while dCas-VP160 may result in an indirect effect by attracting the transcription machinery, which finally generates active epigenetic marks.^43,44^ Interestingly, for dCas9-P300 some sgRNAs were significantly more effective than others, which suggests some particular pattern regarding their position in the gene promoter. In this line, Weissman and colleagues demonstrated that targeted zones where nucleosomes are found impede Cas9 access.^45^ More experiments need to be done in order to confirm this hypothesis.

The absence of synergistic effect when using several sgRNAs with dCas9-P300 was previously reported,^20^ but a selective combination of the most efficient sgRNAs per gene had never been tested. Here, we further demonstrate that sgRNAs that individually may have a significant activation effect, when combined in sets of two or even three, do not necessarily further improve activation. As sgRNA generated *in vitro* could be expensive and laborious, the minor amount of sgRNAs would simplify any application.

Related to potential clinical applications, the proof of concept that it is possible to obtain *beta pancreatic TF* expression using entirely RNA-based CRISPRa could have important practical implications in a near future, considering that RNA delivery is one of the safest systems in the clinical setting.^46,47^ Nevertheless, in our hands, more optimization should be done to improve multiplex gene activation by the entirely RNA delivery method.

Combination of CRISPRa and different culture media conditions, induced *INS* gene expression. Remarkably, NEUROG3 activation resulted in cumulus of insulin granules-like particles in all culture conditions. On the other hand, Ex-4 suplementation allowed *INS* expression in TF -alone or combined-groups. Ex-4, a long-acting glucagon-like peptide-1 (GLP-1) analogue, increases *PDX1* expression by recruiting USF1 and PCAF (histone acetylases) to the proximal promoter in animal models.^48^ The synergistic effect of CRISPRa and Ex-4 could allow sufficient stimuli to induce INS expression and future tests should be performed to address this possibility. Likewise, sequential instead of simultaneous gene activation performed better, which could be expected, because as was noted before, there is a stoichiometrically decreased activation when multiplex stimulation is used.

## CONCLUSION

Among CRISPRa systems tested, dCas9-P300 demonstrated the best benefit/cost-effectiveness for gene activation due to the possibility of reducing the number of sgRNAs to just one or two per gene, while maintaining maximal activation effectiveness. This system allowed multiplex pancreatic beta cell gene activation, in both HEK293 and clinically relevant human iPSCs. Activation of pancreatic lineage transcription factors using CRISPRa resulted in Insulin production, indicating potential to direct differentiation. Finally, an RNA-based system, composed of dCas9-P300 mRNA and synthetic sgRNAs, was able to activate all targeted genes, which may be important for potential clinical applications.

## ACKNOWLEDGMENTS

C.G; L.C; S.R.S, A.T and F.P.B. were financed by CONICET. We thank Marcelo Ielpi and Malena Sosa for help in the cell culture experiments.

## Author contributions

Conceived and designed the experiments: C.G., S.H., R.P.J. and F.P.B.; performed the experiments: C.G., L.C., and F.P.B.; analyzed the data: C.G. and F.P.B.; contributed reagents/materials/analysis tools: S.H. and L.G.; wrote the paper: C.G., RPJ and F.P.B. F.P.B. is the guarantor of this work and as such had full access to all of the data in the study and assumes responsibility for the integrity of the data and the accuracy of the data analysis. All co-authors have reviewed and approved the manuscript prior to submission.

## Authorship Confirmation Statement

The authors declare that this manuscript has been submitted solely to this journal and is not published, in press, or submitted elsewhere.

