## Supplementary material for "Activation of pancreatic β-cell genes by multiplex epigenetic CRISPR-editing": Suplemental Information

Supplementary Table 1. Oligonucleotides used for cloning on sgRNAs expression plasmid (pSPgRNA). Underlined letters indicate overhangs compatible with the BbsI- digested vectors. Bold letters indicate a mismatch in the target sequence (first G allows efficient U6 transcription from expression plasmids).

| Target | Name | Forward Oligo 5´-3´ | Reverse Oligo 5´- 3´ | Reference |
| --- | --- | --- | --- | --- |
| INS (human) | sgRNA1 | CACCGGGGCTGAGGCTGCAATTTC | AAACGAAATTGCAGCCTCAGCCCC | Gimenez, et al, 2016 |
|  | sgRNA2 | CACC**G**CCAGCACCAGGGAAATGGTC | AAACGACCATTTCCCTGGTGCTGG**C** | Gimenez, et al, 2016 |
|  | sgRNA3 | CACC**G**CTAATGACCCGCTGGTCCTG | AAACCAGGACCAGCGGGTCATTAG**C** | Gimenez, et al, 2016 |
|  | sgRNA4 | CACC**G**AGGTCTGGCCACCGGGCCCC | AAACGGGGCCCGGTGGCCAGACCT**C** | Gimenez, et al, 2016 |
| PDX1 (human) | sgRNA1 | CACCGCCCCACGTGGTTCAGCCGG | AAACCCGGCTGAACCACGTGGGGC | Balboa et al., 2015 |
|  | sgRNA2 | CACCGCCTGGCTGGCCGCACTAAG | AAACCTTAGTGCGGCCAGCCAGGC | Balboa et al., 2015 |
|  | sgRNA3 | CACC**G**AGCAGGTGCTCGCGGGTACC | AAACGGTACCCGCGAGCACCTGCT**C** | Balboa et al., 2015 |
|  | sgRNA4 | CACCGTTTGCTGCACACTCCTGAA | AAACTTCAGGAGTGTGCAGCAAAC | Balboa et al., 2015 |
|  | sgRNA5 | CACCGTTTTCGTGAGCGCCCATTT | AAACAAATGGGCGCTCACGAAAAC | Balboa et al., 2015 |
| NGN3  (human) | sgRNA1 | CACCGCACAGCTGGATTCCGGACAA | AAACTTGTCCGGAATCCAGCTGTGC | This paper |
|  | sgRNA2 | CACC**G**CCTCGAGAGAGCAAACAGAG | AAACCTCTGTTTGCTCTCTCGAGG**C** | This paper |
|  | sgRNA3 | CACC**G**TTTGAGGAACCGAGAGTTGC | AAACGCAACTCTCGGTTCCTCAAA**C** | This paper |
|  | sgRNA4 | CACC**G**AGCCTCGTGTGGCTCTGGTC | AAACGACCAGAGCCACACGAGGCT**C** | This paper |
|  | sgRNA5 | CACC**G**CTAGGAGCAAAGCCGTCTG | AAACCAGACGGCTTTGCTCCTAG**C** | This paper |
| PAX4 (human) | sgRNA1 | CACC**G**ATAGCAGATGAAACAGTTGA | AAACTCAACTGTTTCATCTGCTAT**C** | This paper |
|  | sgRNA2 | CACC**G**CAGAGACAGGGGAAGACCTC | AAACGAGGTCTTCCCCTGTCTCTG**C** | This paper |
|  | sgRNA3 | CACC**G**ACCGTCCTGATGCATGCTCC | AAACGGAGCATGCATCAGGACGGT**C** | This paper |
|  | sgRNA4 | CACCGGAGTCTCATCCTTCTGAGG | AAACCCTCAGAAGGATGAGACTCC | This paper |
|  | sgRNA5 | CACC**G**CTGGAAAAGGAGCTCCTAGA | AAACTCTAGGAGCTCCTTTTCCAG**C** | This paper |

| Target | Name | RNA sequence 5' to 3' (target only) | Modification |
| --- | --- | --- | --- |
| PDX1 | hPDX1 sg2syn | GCCUGGCUGGCCGCACUAAG | 2'-O-methyl 3' phosphorothionate in the first and last 3 nucleotides |
| NGN3 | hNGN3 sg2syn | CCUCGAGAGAGCAAACAGAG | 2'-O-methyl 3' phosphorothionate in the first and last 3 nucleotides |
| PAX4 | hPAX4 sg2syn | CAGAGACAGGGGAAGACCUC | 2'-O-methyl 3' phosphorothionate in the first and last 3 nucleotides |
| INS | hINS sg2syn | CCAGCACCAGGGAAAUGGUC | 2'-O-methyl 3' phosphorothionate in the first and last 3 nucleotides |
| INS | hINS sg3syn | CUAAUGACCCGCUGGUCCUG | 2'-O-methyl 3' phosphorothionate in the first and last 3 nucleotides |

Supplementary Table S2. RNA guides as chemically modified RNA.

Supplementary Table S3. Plasmid concentration used in multiplex gene activation related to experiments represented in Figures 2, 4 and 5 in the main text. *The amount corrrespond to each 60mm plate individual group; ** dCas9 expressng HEK293 line, effectors not apply.

|  | MULTIPLEX ACTIVATION | | | | | | | | |
| --- | --- | --- | --- | --- | --- | --- | --- | --- | --- |
| vector/related to | Figure 2 | | | Figure 4 | | | Figure 5 | | |
|  | Individual effector | All effectors | Control | Most active sgRNAs expressing vector | All sgRNAs expressing vector | Control | most active sgRNAs expressing vector | most active synthetic RNA guides | Control |
|  | 60mm plate(ng) | 60mm plate(ng) | 60mm plate(ng) | 12-well (ng) | 12-well (ng) | 12-well (ng) | 12-well (ng) | 12-well (ng) | 12-well (ng) |
| EFFECTOR |  | | |  | | |  | | |
| dCas9-VP160 | 3750* | 1250 | n/a | n/a | n/a | n/a | n/a | n/a | n/a |
| dCas9-TET1 | 3750* | 1250 | n/a | n/a | n/a | n/a | n/a | n/a | n/a |
| dCas9-P300 | 3750* | 1250 | n/a | 750 | 750 | 750 | n/a ** | n/a ** | n/a ** |
| sgRNAs expressing vector |  | | |  | | |  | | |
| *PDX1* sgRNA1 | 52 | 52 | n/a | n/a | 9 | n/a | n/a | n/a | n/a |
| PDX1 sgRNA2 | 52 | 52 | n/a | 45 | 9 | n/a | 225 | 225 | n/a |
| PDX1 sgRNA3 | 52 | 52 | n/a | n/a | 9 | n/a | n/a | n/a | n/a |
| PDX1 sgRNA4 | 52 | 52 | n/a | n/a | 9 | n/a | n/a | n/a | n/a |
| PDX1 sgRNA5 | 52 | 52 | n/a | n/a | 9 | n/a | n/a | n/a | n/a |
| NEUROG3 sgRNA1 | 52 | 52 | n/a | n/a | 9 | n/a | n/a | n/a | n/a |
| NEUROG3 sgRNA2 | 52 | 52 | n/a | 45 | 9 | n/a | 225 | 225 | n/a |
| NEUROG3 sgRNA3 | 52 | 52 | n/a | n/a | 9 | n/a | n/a | n/a | n/a |
| NEUROG3 sgRNA4 | 52 | 52 | n/a | n/a | 9 | n/a | n/a | n/a | n/a |
| NEUROG3 sgRNA5 | 52 | 52 | n/a | n/a | 9 | n/a | n/a | n/a | n/a |
| PAX4 sgRNA1 | 52 | 52 | n/a | n/a | 9 | n/a | n/a | n/a | n/a |
| PAX4 sgRNA2 | 52 | 52 | n/a | 45 | 9 | n/a | 225 | 225 | n/a |
| PAX4 sgRNA3 | 52 | 52 | n/a | n/a | 9 | n/a | n/a | n/a | n/a |
| PAX4 sgRNA4 | 52 | 52 | n/a | n/a | 9 | n/a | n/a | n/a | n/a |
| PAX4 sgRNA5 | 52 | 52 | n/a | n/a | 9 | n/a | n/a | n/a | n/a |
| INS sgRNA1 | 52 | 52 | n/a | n/a | 9 | n/a | n/a | n/a | n/a |
| INS sgRNA2 | 52 | 52 | n/a | 22,5 | 9 | n/a | 112,5 | 112,5 | n/a |
| INS sgRNA3 | 52 | 52 | n/a | 22,5 | 9 | n/a | 112,5 | 112,5 | n/a |
| INS sgRNA4 | 52 | 52 | n/a | n/a | 9 | n/a | n/a | n/a | n/a |
| Empty guide vector | n/a | n/a | 1250 | n/a | n/a | 180 | n/a | n/a | n/a |
| eGFP vector | 250 | 250 | 3738 | 100 | 100 | 100 | 100 | 100 | 1000 |
| Total plasmid mass | 4988 | 4988 | 4988 | 1030 | 1021 | 1030 | 1000 | 1000 | 1000 |

Supplementary Table S4. Primers used for RT-PCR, quantitative PCR (qPCR) , DNA methylation analysis (PCRbis) , ChiP-qPCR and IVT experiments

|  | Forward primer (5’ to 3’) | Reverse primer (5’ to 3’) |
| --- | --- | --- |
| RT-PCR | | |
| *INS* | GCAGCCTTTGTGAACCAACACC | TGTTCCACAATGCCACGCTTC |
| *PDX1* | AAGTCTACCAAAGCTCACGCG | CGTAGGCGCCGCCTGC |
| *NEUROG3* | TCAACTCGGCACTGGACG | AGGGAGAAGCAGAAGGAACAA |
| *PAX4* | GCCACCGGAATCGGACTATC | CCCACGCTGGAACTCTTTCT |
| *BACT* | CCTGGCACCCAGCACAAT | GCCGATCCACACGGAGTACT |
| RT-qPCR | | |
| *INS* | AAGTCTACCAAAGCTCACGCG | CGTAGGCGCCGCCTGC |
| *PDX1* | CGGTAGAAAGGATGACGCCT | TCTCACGGGTCACTTGGACA |
| *NEUROG3* | GCCACCGGAATCGGACTATC | CCCACGCTGGAACTCTTTCT |
| *PAX4* | CCTGGCACCCAGCACAAT | GCCGATCCACACGGAGTACT |
| *BACT* | GGGAGGCGACAAGCGAC | CTGTCGTAGTTCTTCTGGTTTGA |
| dCas9-P300 juntion | GGGAGGCGACAAGCGAC | CTGTCGTAGTTCTTCTGGTTTGA |
| PCR bisulfite | | |
| INS | AATTTAGTTGAGAGAGAAAATTGGA | CCATATACAAACACRCAAAAA |
| ChiP-qPCR | | |
| INS | CGCAGCCTTTGTGAACCAAC | TTCCCCGCACACTAGGTAGA |
| PDX1 | TCTGGCTTGGTTCTTTGTGGT | GATACTCCCGAAATGAGGACG |
| NEUROG3 | TGTCTGTCTGTGATTGGACCG | CCTCCCCCTCGTGTCTTTTG |
| PAX4 | TCAGGAGAGTTGGCGGGTAT | ACGCAAGCTGCTTTCTTTGAG |
| IVT template PCR | | |
| dCas9-P300 | GCATTTAGGTGACACTATAGAAGAGccaagctggctagcgccatg | T(80)tcgaggctgatcagcggttt |

Supplementary Figures

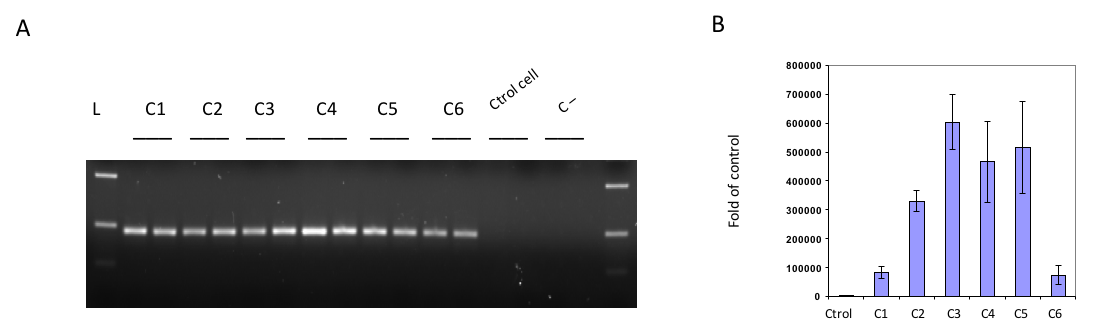

Supplementary Figure S1. DNA integration check in new HEK293 dCas9-P300 expressing line. (A) RT-PCR for dCas9-P300 juntion in six clones. "Ctrol cells" are non-transduced cells, "C-" is PCR control mix. (B) RT-qPCR for dCas9-P300 juntion in the six positive clones to select the clone. Clone number 3 was chosen to expand the line.

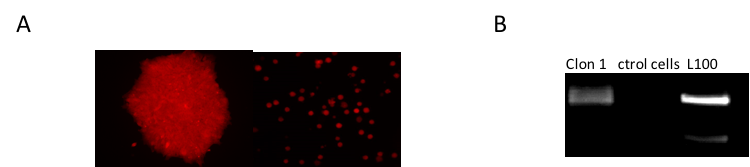

Supplementary Figure S2. DNA integration check in new hiPSC expressing line. (A) Left: Resistent clon human IPS cell (SUN002.1.2 tdt), right: hIPSC non infected (ctrol) (250-400ng/ml puromycin for one week). (B) RT-PCR for dCas9-P300 juntion in positive clon.

***BACT***

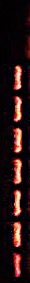

**C+	PDX1	NEUROG3	PAX4	INS	Multiplex Ctrol cells	Ctrol RNA. C-**

_________________________________

**Best sgRNA + dCas9-P300 mRNA in HEK 293 cells**

_______________________

**Individual gene**

Supplementary Figure S3. RT-PCR housekeeping beta actin expression analysis in ordinary HEK293 line transfected with dCas9-P300 mature RNA and the most efficient synthetic sgRNAs. "C+" is human pancreas cDNA; "Individual genes INS, PAX4, NEUROG3, PDX1" are cell groups transfected with the corresponding most efficient sgRNA of each gene; "Multiplex" is the cell group transfected with the most efficient sgRNAs from all genes; "Ctrol cells" is the group without any sgRNA transfected; "Ctrol RNA" is RNA treated by DNAse before cDNA convertion and "C-" is PCR mix control.
